# Survival of prey growing through gape-limited and apex predators

**DOI:** 10.1101/686964

**Authors:** James J. Anderson

## Abstract

A mechanistic model based on first principles of growth and predator-prey behavior introduces the effects of a predator size distribution on the survival of rapidly growing prey. The model, fit to Chinook salmon data, can explain the observed increase in ocean survival with smolt ocean entrance length using different predator size-frequency distributions. The model introduces new dimensions to theories on predator-prey interactions and population recruitment and suggests the possibility that fish recruitment control can be highly variable; sometimes dominated by juvenile growth and encounters with gape-limited predators and other times dominated by adult encounters with apex predators. Additionally, a sensitivity analysis suggests that scale and otolith circuli spacing are insensitive indicators of size-selective mortality but the profile of adult survival with juvenile length provides information on the balance of size-dependent and size-independent mortality processes.

## INTRODUCTION

Understanding survival of fish from juvenile to adult life stages is a major challenge in fishery science and is especially critical to salmon management in which the high variability of spawner recruitment is thought to be strongly influenced by the survival of juveniles during their first few months of ocean residence [1, 2]. Given its importance, early ocean survival has received considerable attention in field studies and reviews. A focal point has been in quantifying the degree of size-selective mortality at different life stages. The rule-of-thumb being that bigger and faster growing juveniles when they enter the ocean in the spring contribute more to the next life stage and ultimately to adult recruitment than do smaller slower growing counterparts [3]. Size-selective mortality is inferred indirectly by back-calculation that compares the otolith and scale circuli patterns of one life stage with the patterns of an earlier stage [4, 5]. A second indirect method correlates the size distribution of cohort members captured at an early life with the population size of adults [6]. These studies and others reveal a variety of patterns, some of which fit in the bigger-faster rule of thumb and others that do not. In support of the rule, several studies have demonstrated yearling chinook salmon and steelhead size and growth during the critical first summer of marine residence, a period to reach a critical size, correlate with later life stage survival and ultimately adult recruitment [5–8]. However, other populations do not fit the critical size and bigger-faster rules. In several subyearling Chinook salmon populations, lower summer growth was associated with higher adult returns [9–11]. One difficulty of size/growth selection involves its magnitude compared to the magnitude of mortality across life stages. In one notable example, a 9% size-selective mortality in British Columbia coho between summer and winter was insufficient to explain the ~70% mortality observed over the period [4]. A follow-up simulation suggested that invoking size as a population control requires an unlikely level of selection directed at the smaller members of a life stage [12]. Adding to the difficulty of early life size-selection are studies that indicate significant ocean mortality on adult salmon [13, 14] and for spawning adults entering their natal river system [15]. In total, the evidence suggests that significant ocean mortality can occur across a range of ages and sizes. However, comparisons of early life size with adult returns of individually tagged salmon clearly indicate that large smolts have higher adult survival [16, 17]. One limitation in resolving the effects of size on fish recruitment has been the lack of a theoretical framework that accounts for the changing spectrum of predators experienced by rapidly growing species like salmon.

This paper presents a mechanistic model that expresses the effects of prey and predator sizes on shaping juvenile to adult survival. The model partitions mortality into size-dependent and size-independent components, which for salmon correspond to gape-limited and apex predators. The two predator classes depend on the predator size-frequency distribution and the predator-prey interactions depend on how quickly the prey grow through the distribution. Fitting the model to data on smolt size vs.adult survival of Snake River salmon reveals that reasonable model fits can be achieved with either predator type dominating mortality. Challenges in identifying the balance of processes and their significance in understanding fish recruitment are discussed.

## MATERIALS AND METHODS

### Model

The model is based on the idea that prey susceptibility to predation decreases as prey progressively grow larger. The prey survival is

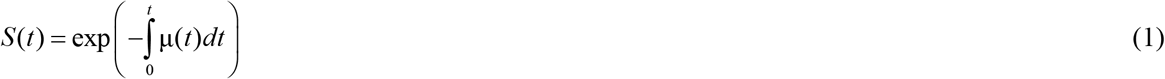

where μ(*t*) is the mortality rate as a function of prey age *t* where *t* = 0 is some reference age when the prey enters the predator field. To incorporate size dependency in the mortality rate, assume predator-prey encounters are Poisson distributed with mean frequency λ and encounter mortality requires the prey size l is less than a random encounter with a predator of gape size, *Z*, i.e. *Z* > *l*. Applying a formulation developed for human survival [18], the mortality rate is 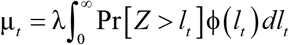 where Pr[*Z* > *l_t_*] is the probability of the predator being larger than prey of size *l_t_* at *t* and φ(*l_t_*) is the distribution of prey lengths at time *t*, which depends on both the prey growth rate and the preferential elimination of smaller prey through predation. There is no closed form for φ(*l_t_*) so applying a discretized approach [19] the mortality rate is tracked for size class increments, *i*, defined by the size of prey at *t* = 0. The class-specific mortality rate is then approximated as μ_*i*_(*t*) = λ Pr[*Z* > *l_i,t_*] and size class-specific survival is

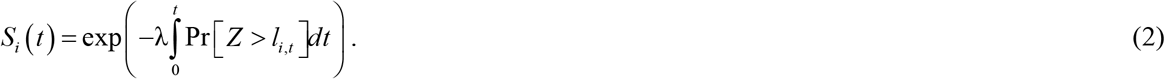

To express the probability that a predator gape is greater than the prey size represent the predator size distribution by a left truncated cumulative normal distribution *F*(*l, m, s,* 0, ∞) where *l* is the prey length, *m* and *s* are the mean and standard deviation of the predator gape distribution. The probability of a predator exceeding prey size class *i* is 1 – *F*, which yields

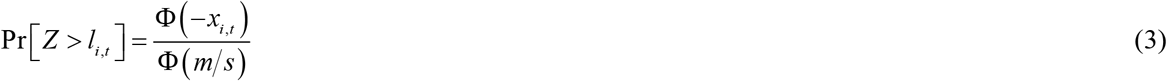

where Φ is the standardized cumulative normal distribution and the normalized prey size is

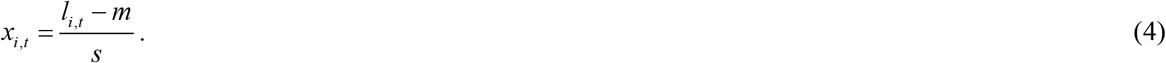

The mortality rate for size class *i* is then

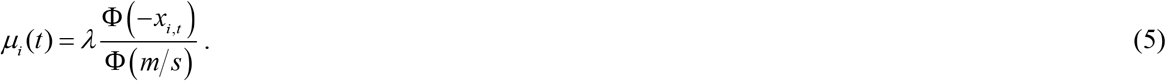

To include age assume linear growth *l_i,t_* = *l*_*i*,0_ + *g_i_ t* where *g_i_* is the growth rate in class *i* and *l*_*i*,0_ is the prey length when entering the predator field at *t* = 0. The normalized prey size as a function of time is

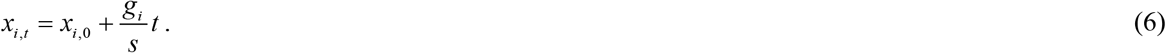

Equation (6) is similar to the early life growth in a Von Bertalanffy growth eqn [20]. While at older ages the equation overestimates size and correspondingly underestimates mortality, the bias has little effect in the model because the majority of size-dependent mortality occurs in early age. Introducing eqns (6) and (3) into (2) the log survival for size class i resulting from length-dependent processes is

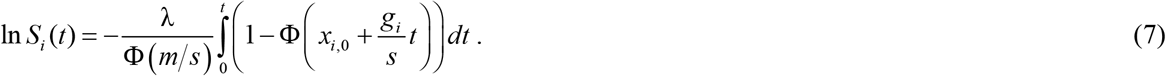

From eqn (6) *s/g_i_ dx* = *dt* and expressing limits of integration in terms of *x*_*i*,0_ and *x_i,t_* then integration of eqn (7) gives the survival as a function of initial size and size at age *t* as

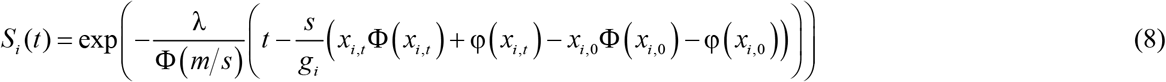

where φ is a standard normalized distribution.

Mortality may also occur independent of the prey length; for example when juvenile [21] or adult [15] prey pass a gauntlet of predators not limited by gape size. This additional mortality can be characterized by a constant mortality rate μ_*H*_ then

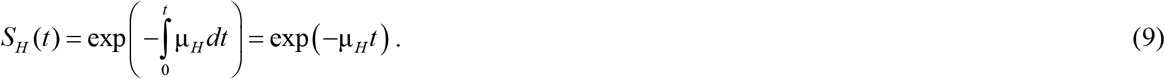

The total survival from length-dependent and length-independent processes for each size class is

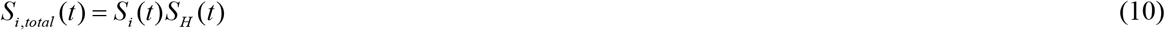

Noting as *t* → that 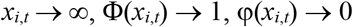, and Φ(–*x*) = 1– Φ(–*x*) then taking the limit of eqn (8) the adult survival for size class *i* is approximated

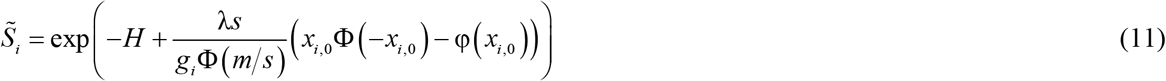

where

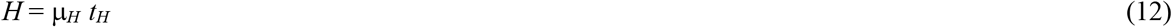

and *t_H_* is the duration of size-independent mortality, which can be assumed to be the age of adult return.

The total mortality rate for the mean prey length is derived from eqns (5) and (12) as

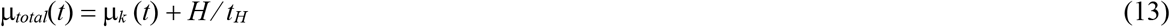

where *k* is the bin index of the mean prey.

To express eqns (8) and (11) in terms of the population average length and growth rate represent size-class growth rate relative to the average growth rate by a logistic curve

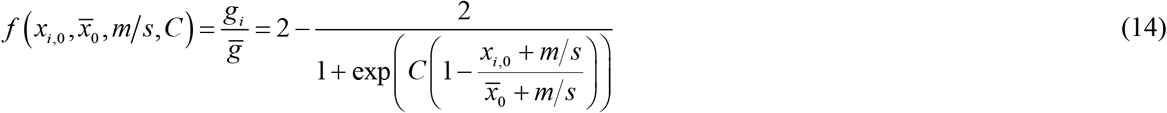

where *x*_*i*,0_ and 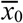 are the normalized prey size for class *i* and the mean size class of the population respectively at time 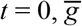 is the mean population growth rate and *C* is a coefficient characterizing the relationship of size class and growth. For *C* = 0 growth rates are equal for all size classes. For *C* < 0 the growth rate is greater than 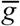 for size classes larger than the mean, which suggests that differences in size at *t* = 0 is the result of different previous growth rates and the pattern is maintained for *t* > 0. Finally, *C* >0 suggests that for *t* > 0 growth rates of the smaller size classes are greater than the mean population growth rate. This condition corresponds to compensatory growth in which prey with slower growth for *t* < 0, as a result of past diet restriction, exhibit catch up growth when the diet restriction is released for *t* > 0 [22, 23].

Note from eqn (4) that the fraction 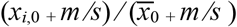 is simply 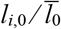 where 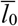 is the mean length of the population at *t* = 0. Also note from eqn (14) that 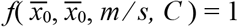.

The final equation for prey survival between predator field entrance and exit becomes

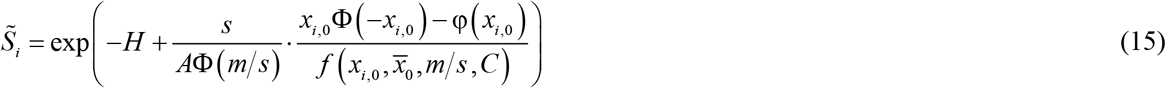

with *x*_*i*,0_ and *x*_0_ normalized fish length defined by the initial fish length *l_i_*, predator gape mean *m* and standard deviation *s*. Each selected *m* and *s* pair defining the predator distribution has associated values of *A, C*, and *H*. The parameter

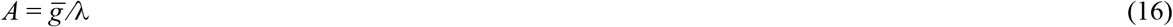

describes the increase in prey length between encounters with gape-limited predators. The parameter *C* is a physiological index of growth at *t* = 0 and H is the total size-independent mortality. From eqns (5) and (16) the ratio of prey population length increase between mortality events with gape-limited predators is

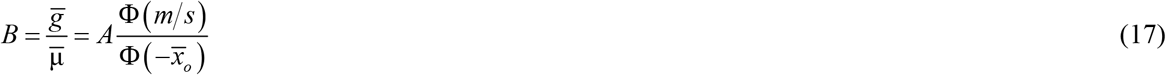

The survival of the mean size class *S*_0_, can be expressed in terms of size-independent survival associated with apex predators *S_apex_* and size-dependent survival associated with gape-limited predators *S_gape_* as

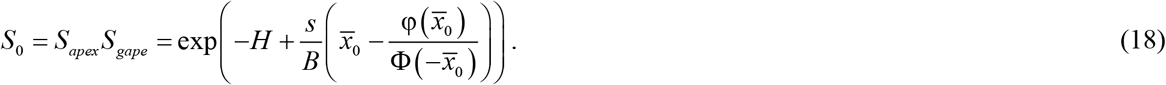

Because the second derivative of eqn (15) is positive for 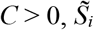 is concave up such that *S*_0_ is less than the mean survival of the cohort. However, difference is small when the degree of concavity is small.

Finally, an index size-selection mortality can be expressed as the ratio of the juvenile mean size of adults to the juvenile population mean size as

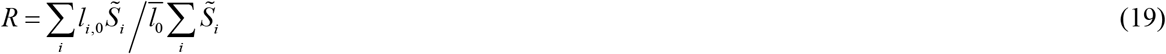

where 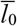 is the mean size distribution of the cohort when entering the predator field, 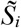 is defined by eqn (15) and *i* = 1, 2, …*I* designates the size classes. Under the assumption of linear growth in early marine residence then *R* also applies to growth circuli spacing in early life stages.

### Estimating parameters

A nonlinear fit of length vs. 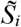 through eqn (15) does not yield a unique estimate of the parameters (*m, s, A, C, H*). However, best estimates of the *A, C, H* triplet do exist for *m, s* pairs and so the parameter space of fitted triplets can be derived by a grid search in which the best triplet is obtained for each *m, s* pair using a nonlinear weighted least-squares algorithm (`nls`) [24]. For each 1 mm incremented *m × s* gridpoint a triplet and its minimum residual square error (RSE) was derived. The *m × s* grid dimensions were determined from predator forging studies discussed below.

### Data

To explore the contributions of size-dependent and size-independent mortality processes in controlling ocean survival the model was fit to data of ocean survival of yearling Chinook salmon (*Oncorhynchus tshawytscha*) vs the fish length prior to ocean entrance. The juvenile fish released as smolts from Snake River hatcheries were collected at Lower Granite Dam, measured for length and tagged with passive integrated transponder tags (PIT tags). Data is available at the DART database (http://www.cbr.washington.edu/dart) and the Pacific states Marine Fisheries Commission (http://www.psmfc.org/). Two years of date were analyzed. Smolts selected for 2008 had been transported through the river in barges and released below Bonneville Dam, the last dam on the Snake-Columbia River hydrosystem. Smolts selected for 2009 had migrated through 7 dams of the hydrosystem. For both passage routes the ocean survival was determined as the smolt-to-adult ratio (SAR), which is the number of returning adults detected divided by the number tagged smolts. The SAR was calculated for 4 mm size increments of smolt migration size. The mean fish lengths for 2008 and 2009 were 136 and 135 mm. These data, discussed in [16], were selected because they represent a factor of 6 difference in SAR as well as the fish experiencing different river passage conditions.

*m, s* – The range of possible predator gape sizes was estimated using the size distributions of prey captured by piscivore predators residing in the Columbia River plume and the Washington coast. Pacific hake (*Merluccius productus*) and jack mackerel (*Trachurus symmetricus*), are major predators and length of their prey from stomach contents range between 50 and 250 mm with a median length of about 150 mm [25]. Avian predators of juvenile salmon in the Columbia River plume and coastal environment, such as Aucklets (*Aethia cristatella*) foraging off Vancouver Island, select salmonids with lengths about 120 mm [26]. At the upper range, apex predators such as salmon sharks have gapes > 800 mm [13]. Clearly, apex predators do not represent the mean gape size but their contribution to predation can be readily represented by grid points with large values of m and s. From these observations the grid range of mean gape size was set at *m* = 50 to 250 mm and the standard deviation of gape size was set at *s* = 4 to 204 mm.

*H* – Size-independent mortality depends on the mortality rate μ_*H*_ and duration *t_H_*, eqn (12). The possible range of *H* varies from 0, corresponding to no size-independent mortality, to the maximum mortality of *H* = –log(*S_crit_*), where *S_crit_* is the adult survival of fish from the largest bin size in a cohort. From, independent estimates of mortality by large gape predators the expected is minimum is *H* = 0.05, which equates with the 4.5% mortality of pinniped predation in the bypass system of Bonneville Dam [15]. Harvest of the Snake River spring Chinook salmon over two decades was under 10% [27] and largely occurred upriver of Bonneville Dam. Thus, harvest mortality was assumed zero. An estimate of *H* can be inferred from a study of ocean survival of Chinook salmon, length 57-100 cm, that were tagged with pop-up satellite archival tags in the Bering Sea and Gulf of Alaska [14]. Salmon sharks (*Lamna ditropis*) accounted for 14 mortalities of the 33 fish with mammals and ectothermic fishes accounting for 2 and 3 fish respectively. A survivorship curve constructed from confirmed predation mortalities indicated survival declined in an exponential-like to 18% over 260 d, resulting in a daily mortality rate of 0. 008 /d. The dominant predator, salmon sharks, reach over 200 cm length [13] and consume Chinook salmon > 80 cm in length. Note that the model fitted ranged of *H* also depends on other model parameters such that under some parameter regions the mortalities associated with apex predators can be subsumed into a size-dependent mortality processes.

The period of size-independent morality *t_H_* again depends on the partition of size-dependent and independent processes. From [13] the period is expected to be on the order of a year or more assuming the Chinook tagged in the [14] study during the winter entered the marine environment in the previous spring. For this analysis *t_H_* is not required in the fitting routine but is required to assess the seasonal pattern of survival according to eqns (10) and (12).

*A* – The ratio of the growth rate to the gape-limited predator encounter rate is derived from eqn (17) using the ratio of growth to mortality rates, *B*, derived from independent rate estimates of mortality 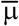 and growth 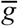. For Snake River spring Chinook salmon, these rates have been estimated over the first few months of marine residence. Growth rates, obtained from otoliths ring widths of Snake River spring Chinook captured off the Washington coast, ranged from 0.5 to 1 mm/d over the first two month of ocean residence [6]. Additionally, using the survival of acoustically tagged juvenile spring Chinook salmon transiting through the Columbia River plume [28], gives a mortality rate of μ ≈ 0.08/d. The estimate is based on a log-linear regression of survival vs. plume residence time and includes an intercept term to account for tagging effects and capture inefficiency of the acoustic detection array. The resulting ratio is 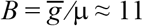. This can then be compared to *B* estimated from eqn (17) for each *m × s* grid point and the corresponding *A* estimate.

*C* – The compensation parameter reflects the differential growth rate of small individuals relative to the mean population growth rate. To estimate the range of this parameter consider laboratory measured compensatory growth in which previously food limited fish exhibited growths 10 to 50% greater than fully fed fish [23, 29]. Assuming that small fish entering the ocean had previously lower food rations than the larger fish, then the range of laboratory observed growth rates correspond to *C* values between 0 and 3 (Fig. 1).

**Figure 1.**
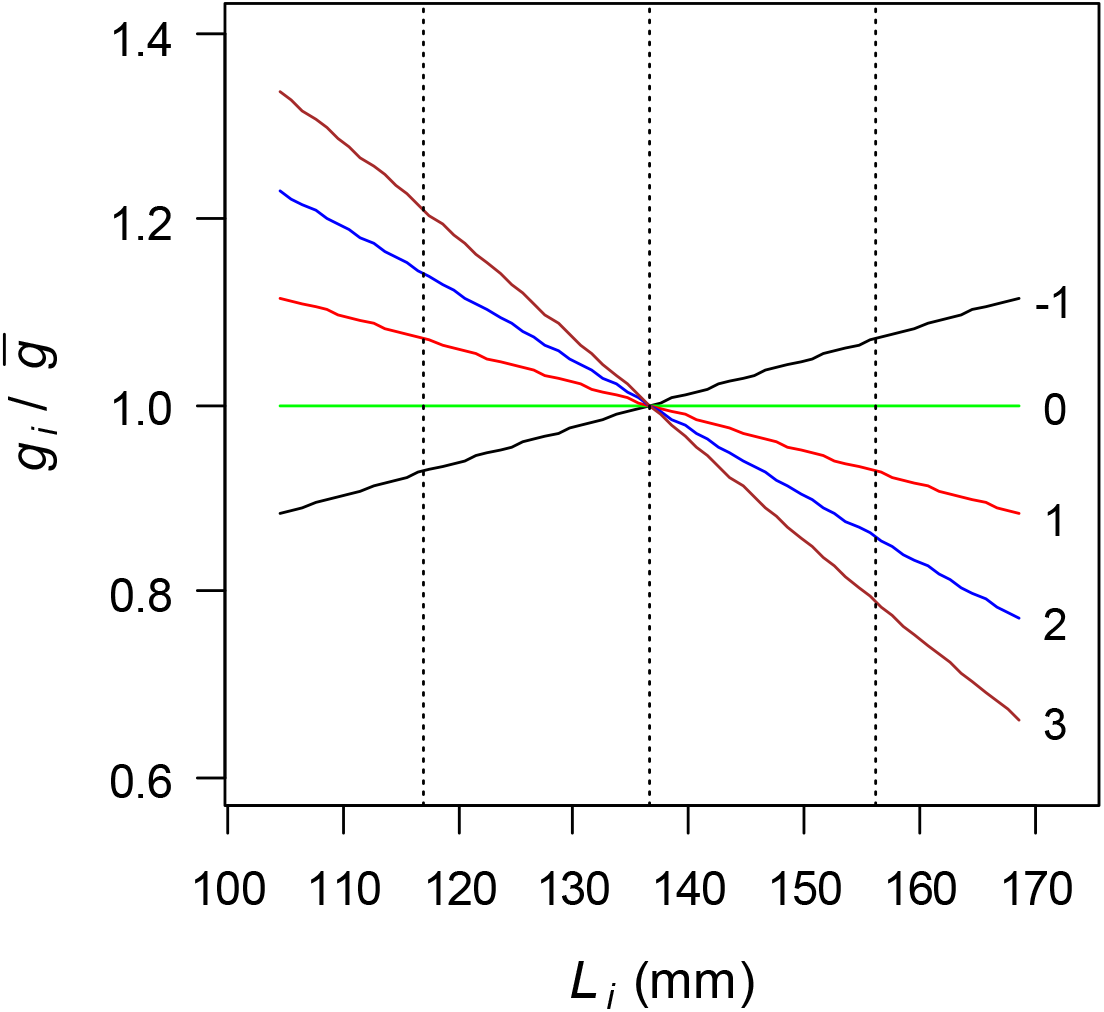
Effect of fish length on relative growth rate, equation (14) for values of C

## RESULTS

Figure 2 illustrates the parameter space depicted as individual plots of *m, s* grid points and corresponding fitted triplets (*A, C, H*) against the minimum RSE value for each point. The figure also includes *B* points calculated from eqn (17). The important ecological features of the parameter space can be viewed in the context of two regions in the *H* vs RSE plots. Region I consists of a horizontal cloud of points demarking the upper boundary of *H* and Region II consists of the lower boundary of *H*. Note each *H* point has corresponding points in the other plots of Fig. 2, which together generate a corresponding SAR vs length curve. Region I contains parameters associated with high size-independent mortality and Region II contains parameters associated low size-independent mortality. The vertical columns of *H* points clustered around RSE = 0.19 for 2008 and 0.065 for 2009 depict the transition region of fitted solutions between the boundary regions. While a continuum of parameter combinations fits the SAR vs length patterns, the basic feature of the patterns is illustrated by considering the minimum RSE points from the two boundary regions (Table 1).

**Figure 2.**
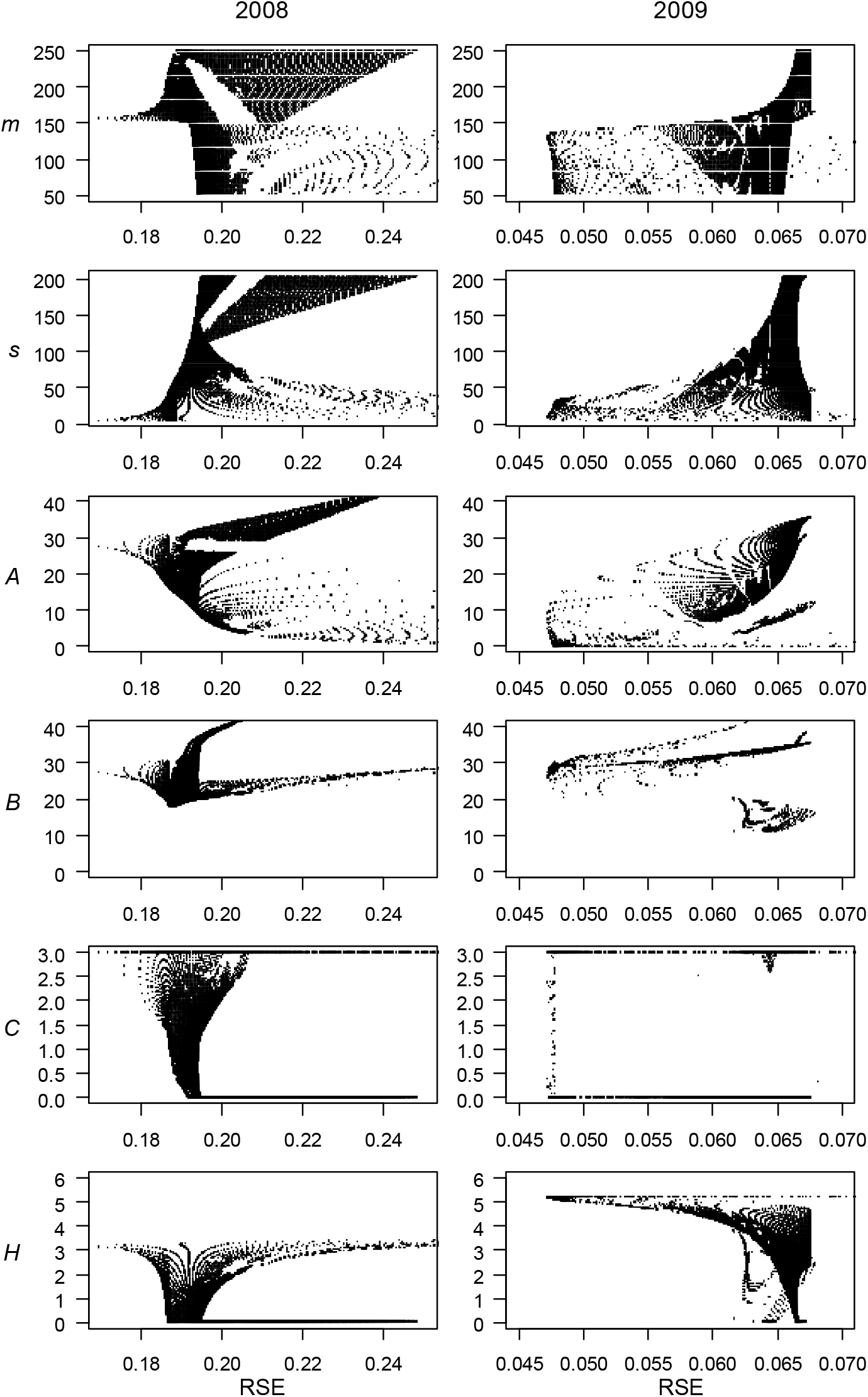
Parameter space vs RSE for years 2008 and 2009. Points in *A, C* and *H* figures represent best fits for *m × s* grid points. Points in *B* figures are calculated from eqn (17). Regions are classified by the *H* vs RSE plots. Region I points encompass the upper boundary of H and Region II points encompass the lower boundary. The transition region corresponds to the vertical columns of points between the two boundaries

**Table 1.**
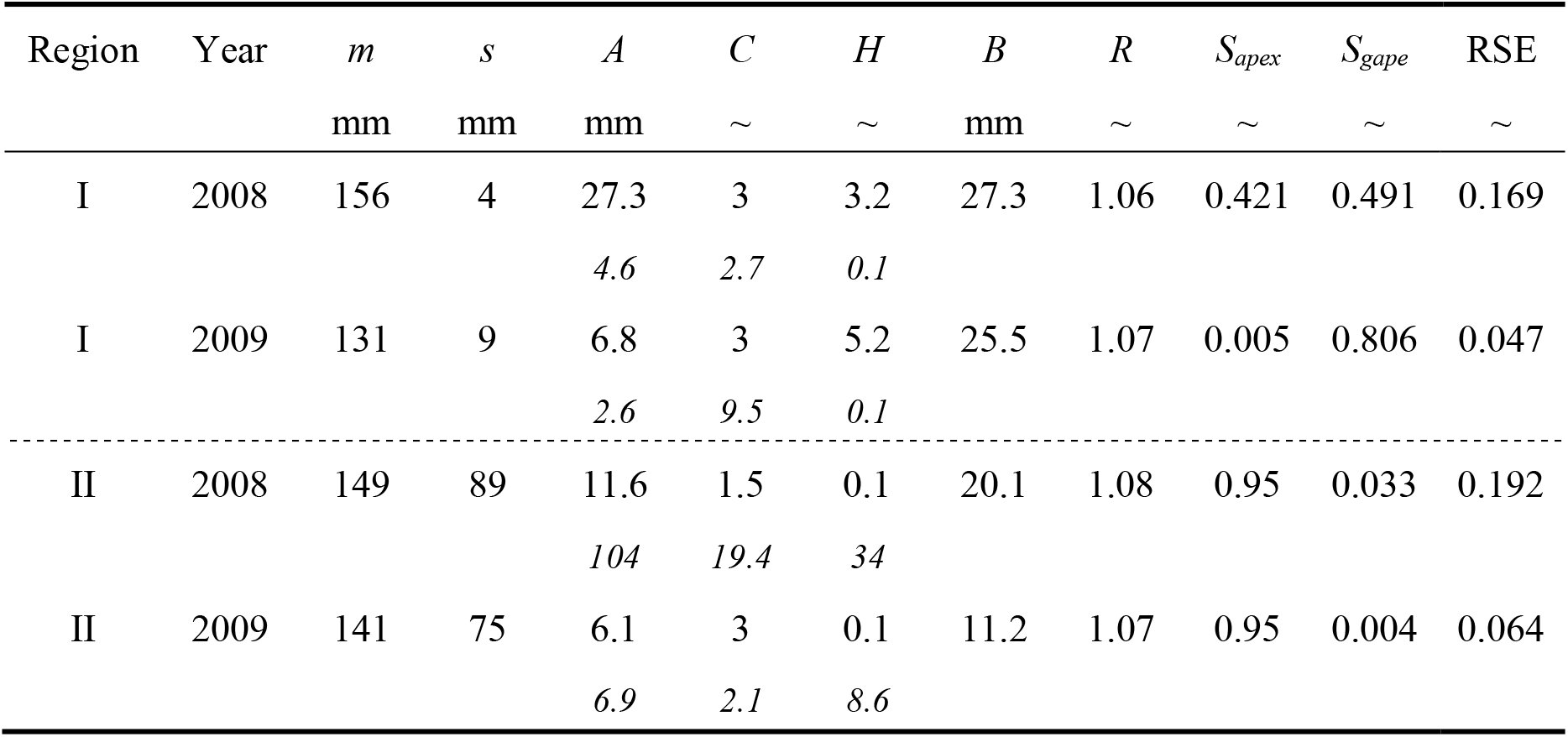
Parameters for Snake River spring Chinook salmon migration year 2008 and 2009 for two regions of the parameter space. Italic numbers depict standard errors from nls fitting

For both Regions and years, the mean gape values are similar, *m* ≈ 144 ± 11 mm, however the gape std, *s*, is small in Region I and large in Region II (Table 1). This pattern is further illustrated in Fig. 3, showing an interpolated contour of *H* on the *m x s* surface. As gape mean and std increase size-independent mortality, *H* decreases because the fit is obtained with more size-dependent predators and fewer size-independent predators. Also important, the growth to mortality rates, *B*, are similar for the two years in Region I but different in Region II. In contrast, *R*, corresponding to the strength of size selection mortality, exhibits little variability across years and Regions.

**Figure 3.**
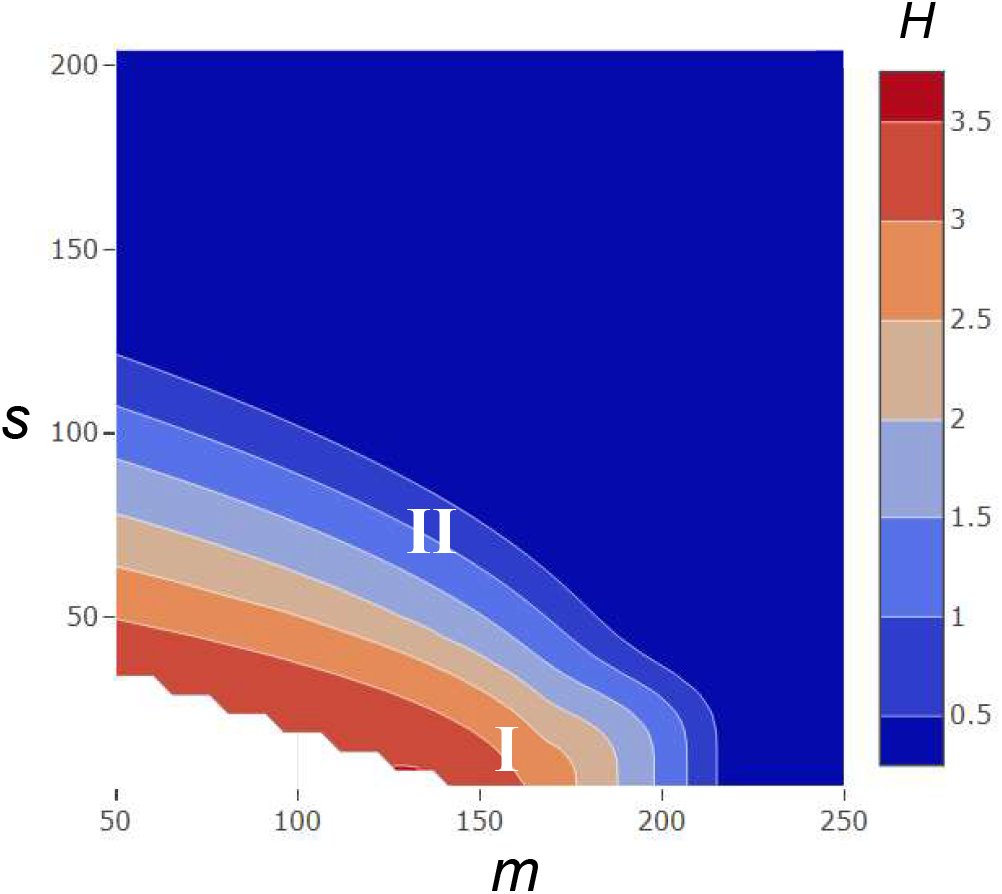
Contour plot of *H* on a *m × s* surface for 2008 migration year. Contours produced with plotply program in R^©^. Parameter Regions corresponding to Table I depicted by I and II.

Figure 4a illustrates the fit of SAR vs *l* for Region I, which captures a distinct step-like pattern for both 2008 and 2009. The normalized gape distribution (Fig. 4b) illustrates the mechanism behind the step. For 2008, all but the largest fish enter the ocean smaller than the mean gape size, which results in greater size-dependent exposure to predators and greater curvature in the SAR vs *l* profile, which only flattens for the largest fish (Fig. 4a). In contrast, for 2009 half the fish enter the ocean larger than the mean gape (Fig. 4b) and therefore the SAR vs *l* distribution flattens quickly for fish larger than the mean (Fig. 4a). For Region II parameters the predator gape distributions for the two years are similar (Fig. 5b), the curvatures of the SAR vs *l* profiles are similar and neither year exhibits a step pattern (Fig. 5a).

**Figure 4.**
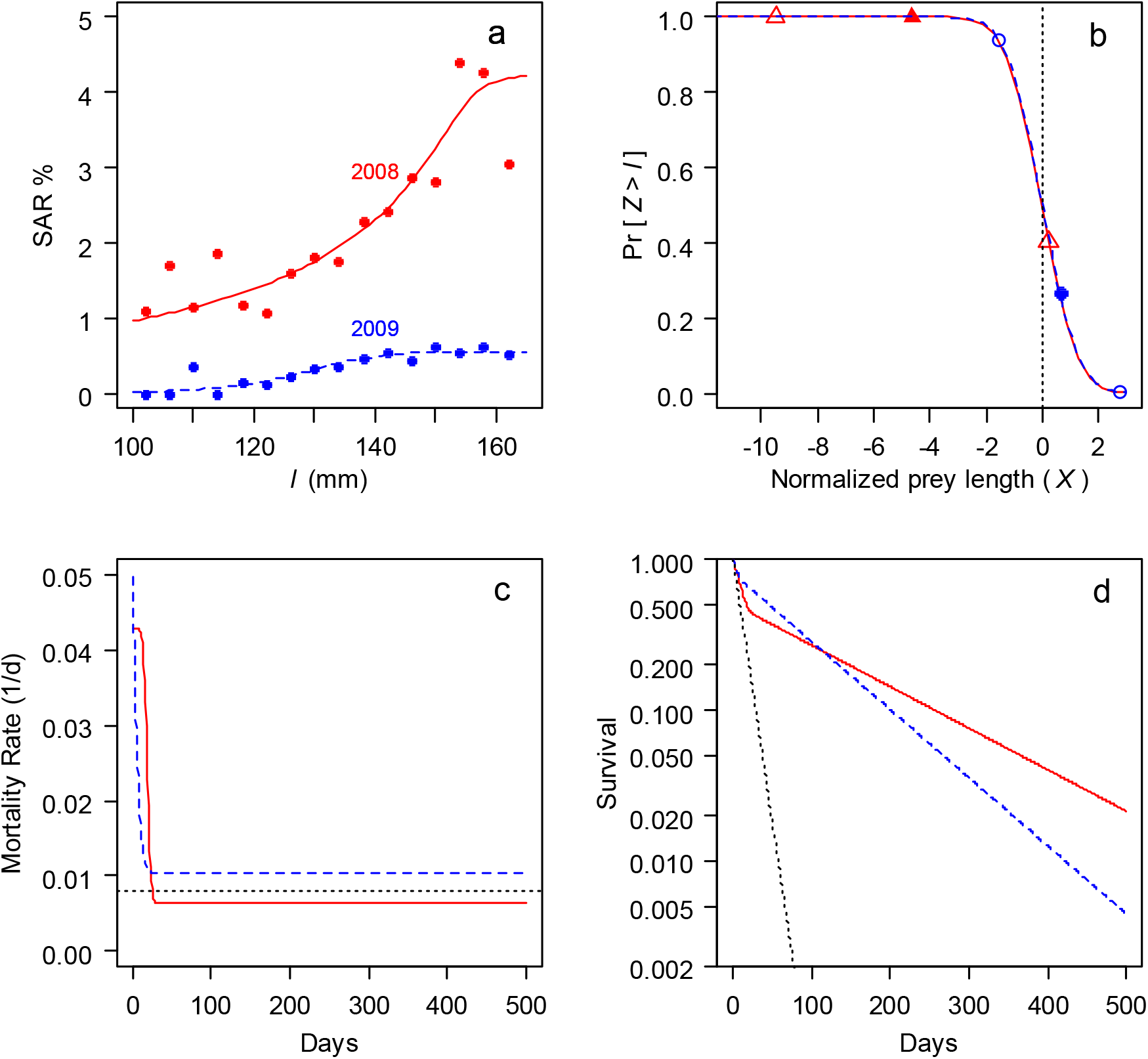
Model fits using Region I of parameter space for Chinook salmon migration years 2008 (red solid lines) and 2009 (blue dashed lines) (Table 1). (a) SAR vs juvenile fish length binned in 4 mm intervals with lined produced by eqn (11) using parameters from Table 1. (b) Probability that predator length is greater than prey length in units normalized by eqn (4) with year 2008 mean 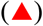 and std 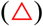 and year 2009 mean 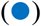 and std 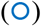. (c) Mortality rate from eqn (13) with dotted line showing apex predator rate from [14] assuming *t_H_* = 500 d. (d) Survivals for 2008 and 2009 from eqn (10). Dotted line is survival estimated from [28].

**Figure 5.**
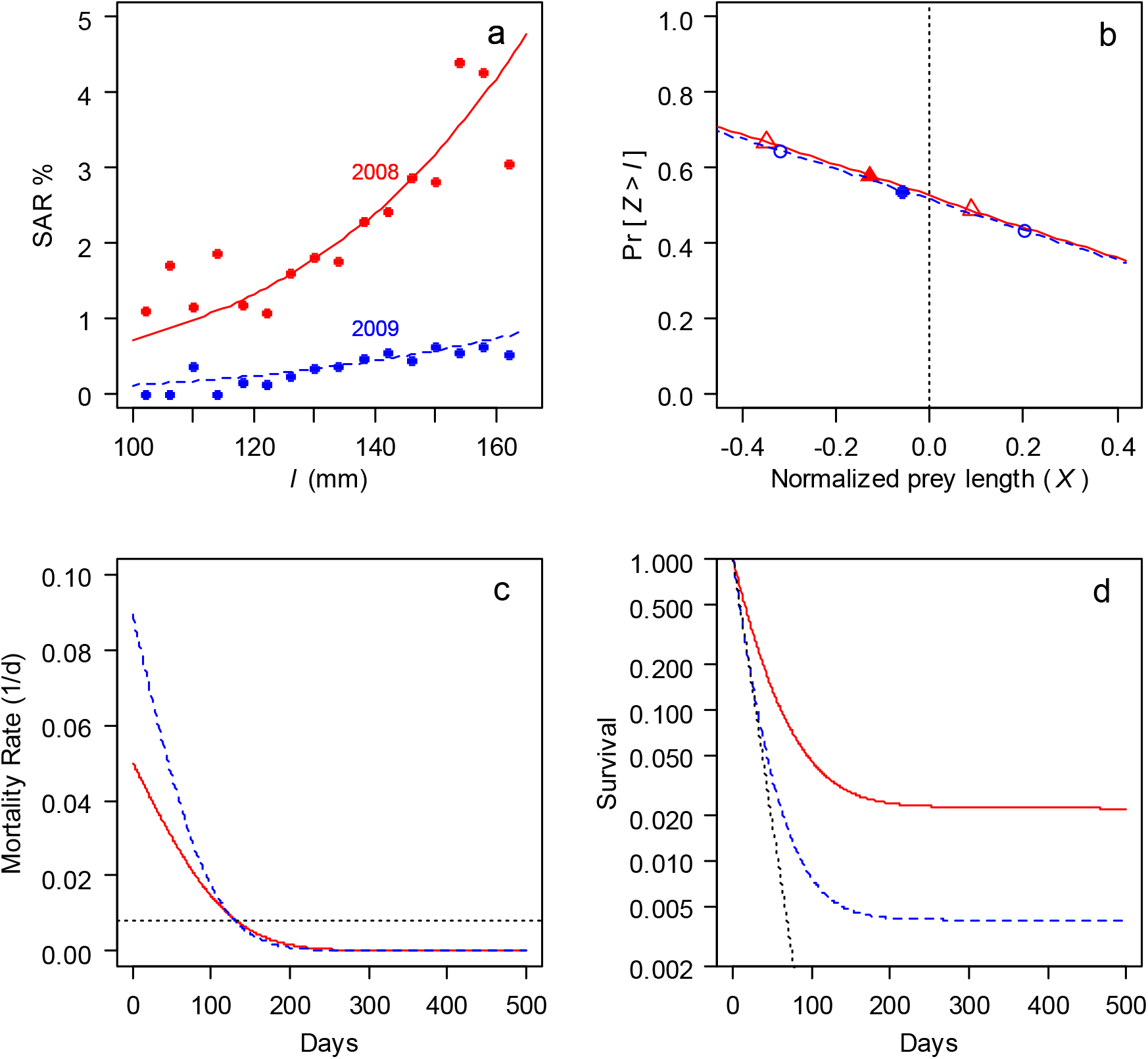
Model fits using Region II of parameter space for Chinook salmon migration years 2008 (red solid lines) and 2009 (blue dashed lines) (Table 1). (a) SAR vs juvenile fish length binned in 4 mm intervals with lined produced by eqn (11) using parameters from Table 1. (b) Probability that predator length is greater than prey length in units normalized by eqn (4) with year 2008 mean 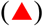 and std 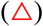 and year 2009 mean 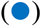 and std 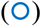. (c) Mortality rate from eqn (13) with dotted line showing apex predator rate from [14] assuming *t_H_* = 500 d. (d) Survivals for 2008 and 2009 from eqn (10). Dotted line is survival estimated from [28].

The temporal differences between Region I and II are demonstrated by the mortality rate (Figs. 4c, 5c) and survival profiles (Figs. 4d and 5d). For Region I, the mortality rate declines within the first month of ocean entrance as fish grow larger than the narrow size distribution of gape-limited predators (Table 1). As a result, the duration of size-dependent is brief and size-independent mortality dominates during ocean residence (Fig. 4c). For 2008 and 2009 the size-independent mortality rates bracket the apex predator rate estimated from [14]. This pattern of mortality produces a rapid decline in survival over the first month of ocean residence followed by a slower exponential decline (Fig. 4d) until *t* > *t_H_* where *t_H_* is either the age of maturation or the age at which fish escape apex predators. Additionally, for Region I parameters the difference in SAR between years is largely due to the difference in encounter frequencies with apex predators as quantified by exp(*H*_2009_ /*H*_2008_) = 5.1. This difference approximates the ratio of SARs of the largest bins for the two years. For Region II, the size-dependent mortality rate gradually declines as fish grow through the wider distribution of predator gapes (Fig. 5c). In this case, the initial size-independent mortality rate at ocean entrance is similar to that calculated from acoustically tagged fish in the Columbia River plume, ~ 0.08 d^-1^ [28] and the rate at ≈ 120 d equals that of apex predators [14]. Note, the lower limit of size-independent mortality rate was set to pinniped predation rate [15]. Correspondingly, with a low mortality rate at older ages the survival asymptotically approaches the mean SAR (Fig. 5d).

## DISCUSSION

The model formulates effects of fish growth on exposure to gape-limited and apex predators from first principles of growth, expressed by 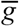 and *C*, the predator size distribution, expressed by m and s and encounters with gape-limited and apex predators, expressed by λ and μ_*H*_ Fitting the model to survival vs juvenile length for a given predator distribution yields the growth parameter *C*, the ratio of growth to predator encounter frequency *A*, and the apex predator mortality index *H*. The effects of prey growth and size on survival is then characterized by *m, s, A, C* and *H*, which can be estimated independently.

The model properties are illustrated using SAR vs ocean entrance length data of Chinook salmon from the Snake River of the Pacific Northwest. The model fit to the data reveals processes controlling marine survival that can be explained by alternative ecologies, which are represented by two different regions of the model parameter space. In Region I, gape-limited predators, such as piscivorous birds and fishes, exhibit a narrow gape-frequency distribution resulting in the majority of predation ascribed to apex predators with gapes exceeding the adult salmon size. The Region II predator size distribution is wider resulting in a more diverse population of gape-limited predators. With Region I ecology, fish recruitment is largely insensitive to ocean growth while with Region II ecology recruitment depends on growth over the first year or more of ocean residence. For the two years in which the model was tested Region I parameters best fit the SAR vs length data, suggesting the importance of apex predators in controlling recruitment. However, both regions capture estimated mortality rates for early and later ocean residence.

The results demonstrate the importance of the predator size distribution in controlling stock recruitment and thus should deepen and extend current theory. Firstly, the model introduces predator size into the critical period hypothesis in which early growth determines recruitment [3]. The original hypothesis was based on the observation that coho salmon with higher spring growth rates were more likely to survive the winter than counterparts with lower rates. This hypothesis comports with Region II parameters in which a wide distribution of predator gapes results in growth controlling survival over several months. Thus, the critical period hypothesis might be most applicable in Region II regimes resulting in smooth SAR vs length and time profiles as depicted in Fig. 5. In contrast, fish entering a Region I environment, as depicted by Fig. 4, would encounter predators with a small gape range resulting in step-like SAR vs length profiles and importantly early ocean growth would have a weak influence on recruitment. Thus, the strength of critical period growth on recruitment might require a wide predator gape distribution that could be revealed by a smooth SAR vs length relationship. In addition, the circuli spacing of the coho survivors, which from eqn (19) equates to *R* = 1.1, is suggestive of a Region II environment in which R values are slightly larger (Table 1). However, a sensitivity analysis over the parameter space indicated that *R* changes little, suggesting circuli spacing is an insensitive indicator of population control processes.

The model also provides a perspective to the balance of top-down vs bottom-up control of recruitment, i.e. the TB balance. Here, the predator size distribution partitions the TB balance across short and long term scales. Over the short term, the ratio of growth to gape-limited predation, B, characterizes the TB balance. Using the Snake River Chinook salmon in which *B*_2008_ > *B*_2009_, then with Region II parameters growth is a more important factor than predation in 2008 than in 2009 assuming the mortality rate is similar between years. However, with Region I parameters the balance is the same for both years (Table 1). Over the long term the top-down processes dominate if *S_gape_* > *S_apex_* and bottom-up processes dominate, i.e. growth becomes a factor in survival, if *S_gape_* < *S_apex_*. In general, top-down control corresponds to systems with Region I parameters and bottom-up control largely corresponds to Region II parameters. Thus, in the model the TB balance can shift in time depending on the preys’ initial size and growth plus the predators’ density and size distribution. For the two years under Region I parameters, top-down processes control recruitment in 2009 and bottom-up processes control recruitment in 2008. However, under Region II parameters bottom-up processes control recruitment in both years (Table 1). Thus, the TB balance may vary interannually.

The model provides information on the value of circuli growth increments as indicators of size-selective mortality. The change in the early growth patterns from the early population to the survivors has been hypothesized to reflect the strength of size selection on survival [3]. However, mortality associated with stage changes is smaller than the actual mortality across the stages [e.g. 11, 12]. The model suggests a possible reason for this discrepancy. Over the parameter space *R* varies between 1.0 and 1.10 with mean value of 1.07 ±0.1 (Table 1). Thus, *R* changes little suggesting that circuli width is not a sensitive indicator of the strength of size-selective mortality.

The model derives population outcomes with two fundamental assumptions that require discussion. The first assumption is that the predator-prey spatial-temporal interactions can be represented in a prey reference frame. The justification is based on the premise that predators exhibit ideal free foraging [30] in which the predator habitat coincides with the temporal-spatial patterns of prey smaller than the predators’ gape. Thus, escaping predation through growth is equivalent to escaping predation by exiting the predator habitat. The second assumption is linear growth, which is ultimately violated because growth rate declines with age. However, the effect of overestimated size at older ages is mitigated by the declining contribution of mortality with increasing prey size.

In summary, the model introduces the effects of the predator size distribution on the survival of growing prey. This link is not easily incorporated in current predator-prey models and so these models are arguably incomplete for generalizing across different predator fields. The model presented in this paper has few parameters and readily fits to salmon data illustrating that the predator size distribution is critical in shaping the time course of survival. In particular, the analysis comports with recent Chinook salmon predation studies and suggests that recruitment is not wholly controlled by processes occurring during early ocean residence. However, the degree of early and late control of salmon recruitment is not resolvable in the limited analysis of this paper. Fortunately, analysis of existing data on SAR vs fish size should provide more information on the issue. However, advancing the understanding of recruitment and more fundamentally the processes controlling survival of rapidly growing prey will need further information on both the size distributions of prey and their predators as well as the frequency and outcomes of their interactions. The methods exist to obtain such data [e.g., 28, 31] but the resources needed are significant.

## Acknowledgements.

I wish to thank Jeff Rudder and Jennifer Gosselin for helpful discussion on the model development. Study was supported by Bonneville Power Administration contract 76910 REL 04. The author has no conflict of interest.

